# Automatic detection of nociceptive pain levels using frequency bands from electroencephalographic (EEG) signals

**DOI:** 10.1101/2025.03.05.641582

**Authors:** Rogelio Sotero Reyes-Galaviz, Luis Villaseñor-Pineda, Camilo E. Valderrama

## Abstract

Pain is considered an unpleasant but vital experience for every living being, it is extremely complex and subjective because it is composed of different variables related to their experiences. Their history, biological sex, socio/cultural context, mood, and hormonal changes can affect their perception. Nociceptive pain is more linked to tissue damage or stimulus, and this will always have a reaction that goes from its activation in the nociceptors in the peripheral nerves of the living being to the central nervous system. The most common way to assess pain today is to apply numerical scales or ”How much pain do you feel?” questionnaires, which are usually falsifiable and unreliable. Therefore, this work seeks to make use of biosignals such as Electroencephalography (EEG) to identify and evaluate pain at different levels. This nociceptive pain is generated by applying a laser on the back of the hand, which consists of three different intensities. It has been possible to differentiate between two levels of pain (high pain and low pain) with 83% accuracy using information from the power of frequency bands of the brain signal. Indicating that there are differences in the powers of the frequency bands as pain increases.

**Author summary:** **Rogelio Sotero Reyes-Galaviz:** Mechatronic engineer from the Polytechnic University of Tlaxcala (UPTlax), with a Master of Science degree in Biomedical Science and Technology at the Instituto Nacional de Astrofísica, Óptica y Electrónica (INAOE). He is currently pursuing a PhD in Biomedical Sciences and Technologies at INAOE. His research is focused on pain quantification using electrical brain signals (EEG) and machine learning methods. His lines of interest are Signal Processing, Electroencephalography, Stress, Music Therapy and Pain. **Luis Villaseñor-Pineda:** Luis Villaseñor received his PhD degree in Computational Sciences from l’Université Joseph Fourier (now Université Grenoble-Alpes), France, in 1999. He is currently a senior researcher in the Computational Sciences department at the Instituto Nacional de Astrofísica, Óptica y Electŕonica, México, and a member of the Mexican Academy of Sciences, the Mexican Association of Natural Language Processing, and the Mexican System of Researchers (Level II). His research interests focus on human-computer communication using human language as well as different biosignals (speech, brain-signal, etc.).

**Camilo E. Valderrama:** Assistant Professor in the Applied Computer Science department at the University of Winnipeg, specializing in the application of machine learning, statistical models, and signal processing to extract meaningful patterns and support decision-making processes. His research spans diverse areas, including affordable fetal monitoring, reducing redundant laboratory tests in intensive care units, protecting children from unhealthy-food advertising, and validating neuromarketing principles. Prior to this, he completed a two-year postdoctoral fellowship at the University of Calgary. He holds a Ph.D. in Computer Science with a concentration in Biomedical Informatics from Emory University (Atlanta, GA, USA), a Master of Science in Informatics, and a Bachelor of Science in Software Systems Engineering from Universidad Icesi (Cali, Colombia).

## Introduction

Pain is such a complex experience that it can be compared to love, only those who have loved before can empathize with the other. This happens with pain, only those who have experienced it can imagine the unpleasant suffering that the other is undergoing. And this experience is so inherent to the living being because it has been evolving along with us, adapting and perfecting itself. History demonstrates that pain has accompanied multicellular living beings with vestiges of nerve fibers since ancient times throughout their evolution and has conditioned them to become the beings we know today. There is evidence from living beings in the time of the caves which can be seen with fractures, dental pain, bone tuberculosis, inflammatory processes, and even childbirth that allows us to know how this has accompanied and punished all since those days.

Pain is such a frequent symptom in our days that it seems like an ally in some cases, warning us of possible diseases or serious ailments as a natural early diagnosis. On the other hand, when perceiving it with greater frequency or in greater intensities, its perception can be complicated by mixing with different emotional and sensory variables that make this response greater than the physiological disease itself. The logical way to think about its perception is that all living beings have a pain threshold that will respond similarly to equal stimuli. Still, the behavior and emotional part of pain completely modify everything, making it so difficult to understand, quantify, and treat. Since when talking about pain, what the affected person feels is underestimated or overestimated, nowadays the evaluation of pain depends a lot on the final judgment of a professional. Accompanying their verdict, different techniques have been created to quantify pain. Among some of them are the analogical scales in which the person has to say in levels how much pain they feel, these can be verbal (low, medium, high), numerical (0 - 100), or visual analog (mark on a line from 0 to 10 cm). Others have been created for people with little schooling where depending on the size of objects, for example, fruits from grapes to watermelons, they must say the intensity of the pain. Other times patients are asked to draw where and how it hurts, or a series of questionnaires are made to know the disabilities generated by the pain or the adjectives that best describe it. Of the above, some turn out to be better or worse for certain populations, but it is still the way to “quantify” pain most used by all doctors. However, they are still unreliable and useless for people with communication problems, since direct interaction with the affected person is needed.

As a result, there has been a growing interest in going beyond understanding, identifying, and quantifying it. This has been done using biosignals. Since the pain pathway from the emitting organ or tissue to the central nervous system is known in some way, some data capture techniques can be used in some areas of interest related to this experience and work with it. In the case of this research, Electroencephalography (EEG) is being used, a technique where a series of sensors are placed on the scalp to obtain the brain’s electrical response caused by the interaction of neurons at all times, and more so when there is a stimulus so powerful such as pain [1–3]. The database was recorded by Tiemann et al. [4] in 2018, and now is public. They recorded the EEG of people under nociceptive pain, which is related to tissue damage or stimulation that activates fibers called nociceptors, and then the pain pathway starts, as can be seen in Fig 1. These reactions were caused by a transcutaneous laser applied at 3 different intensities per subject. In their work is hypothesized that each intensity will generate a type of pain. They found that there is a series of patterns in the EEG signal at the moment of pain, it generates certain peaks of the signal which have to serve as a way to detect it, but not to quantify it.

**Fig 1.**
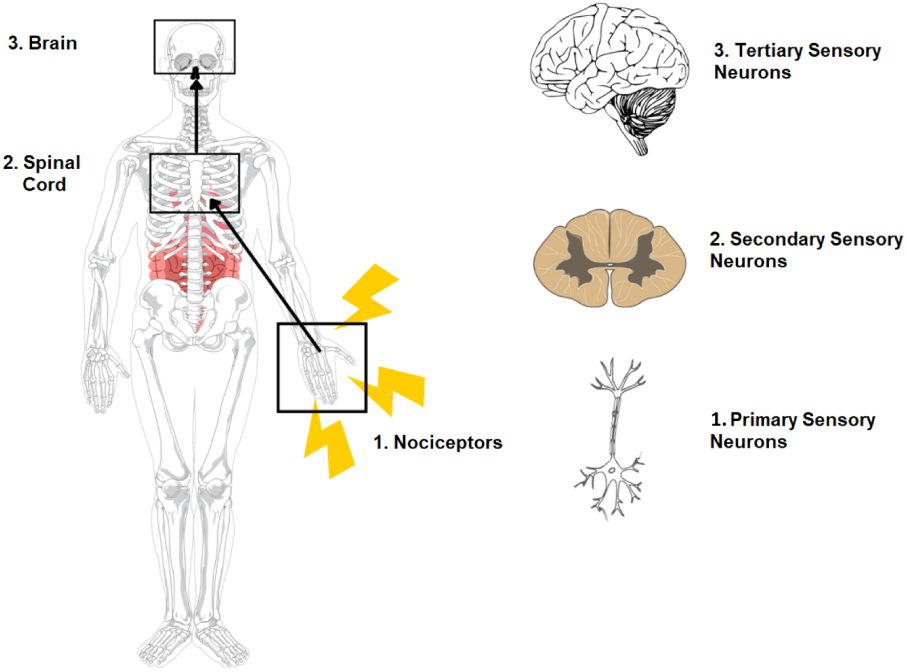
Nociceptive process pathway in humans divided into three blocks. Starting with the Nociceptors, then the Spinal Cord, and finally the Brain.

This paper presents an approach to predicting pain levels using signal processing and machine learning techniques. The paper analyzes pain detection considering several aspects. First, it addresses the discrimination, based on EEG signals, of different pain levels. A preliminary experiment seeks to verify whether sufficient information exists to distinguish between the pre-stimulus (moments before the application of the stimulus) and nociceptive pain. Second, it examines the discrimination between different pain levels. Two scenarios were designed: one where the pain levels are differentiated (high and low), and another, more complex scenario with a continuum of three pain levels. Two approaches were used to determine the labels corresponding to each pain level. Finally, the contribution of the EEG channels was evaluated, using either the information from all channels (62) or focusing exclusively on the channels associated with the somatosensory cortex (20).

### Related work

Assessing pain through electroencephalography (EEG) has gained traction as a promising method for objectively measuring the perception and its underlying neural mechanisms. One of the key advantages of using EEG is its ability to provide real-time data on brain activity in response to painful stimuli. For instance, studies have shown that EEG changes can occur even when traditional pain assessments, such as thermal pain tests, do not indicate significant differences between groups, suggesting that EEG may be more sensitive to the underlying pain mechanisms [5]. This sensitivity is particularly relevant in pediatric populations, where chronic musculoskeletal pain can manifest differently than in adults.

Furthermore, specific EEG patterns, such as alterations in alpha band activity, have been correlated with pain severity, indicating that these oscillations may serve as biomarkers for pain perception [6–8]. Other studies have also explored other frequency bands such as beta [2, 9, 10], or identified certain brain regions that are activated more frequently when pain is present (precentral gyrus, postcentral gyrus, anterior cingulate cortex, and insula) [11]. Additionally, some researchers have attempted to classify pain levels. For instance, Elsayed [12] reported achieving 94.4% accuracy in distinguishing between three levels of pain.

Significant challenges remain in this field, primarily due to the subjective nature of pain and its variability across individuals. The central goal is to identify objective markers through the analysis of biosignals —EEG signals in this work— that enable a more reliable assessment of pain presence and intensity, reducing reliance on patient self-reporting.

## Materials and methods

The database used in this work was published in 2018 by Tiemann et al. [4]. Everything was recorded using the Brain Vision Recorder (Brain products) software and Brain Amp Plus electroencephalogram. In their study, 51 right-handed and completely healthy participants (25 women and 26 men) attended 4 sessions, with an average age of 27 years (20 - 37). Everything was approved by the local ethics committees, and the participants were informed about the procedure. The study protocol is detailed below.

### Experimental protocol

The experiment consisted of 3 core conditions (perception, motor, and autonomic). In addition, a fourth condition was recorded, where each core condition was present and mixed, as seen in Fig. 2. The four conditions were applied to each participant. In each session, 60 applications of the laser stimulus were applied to the back of the left hand. The laser intensity varied from low, medium, and high, each one repeated 20 times in a pseudorandom manner. Between each application, there was a resting time of 8 to 12 seconds.

**Fig 2.**
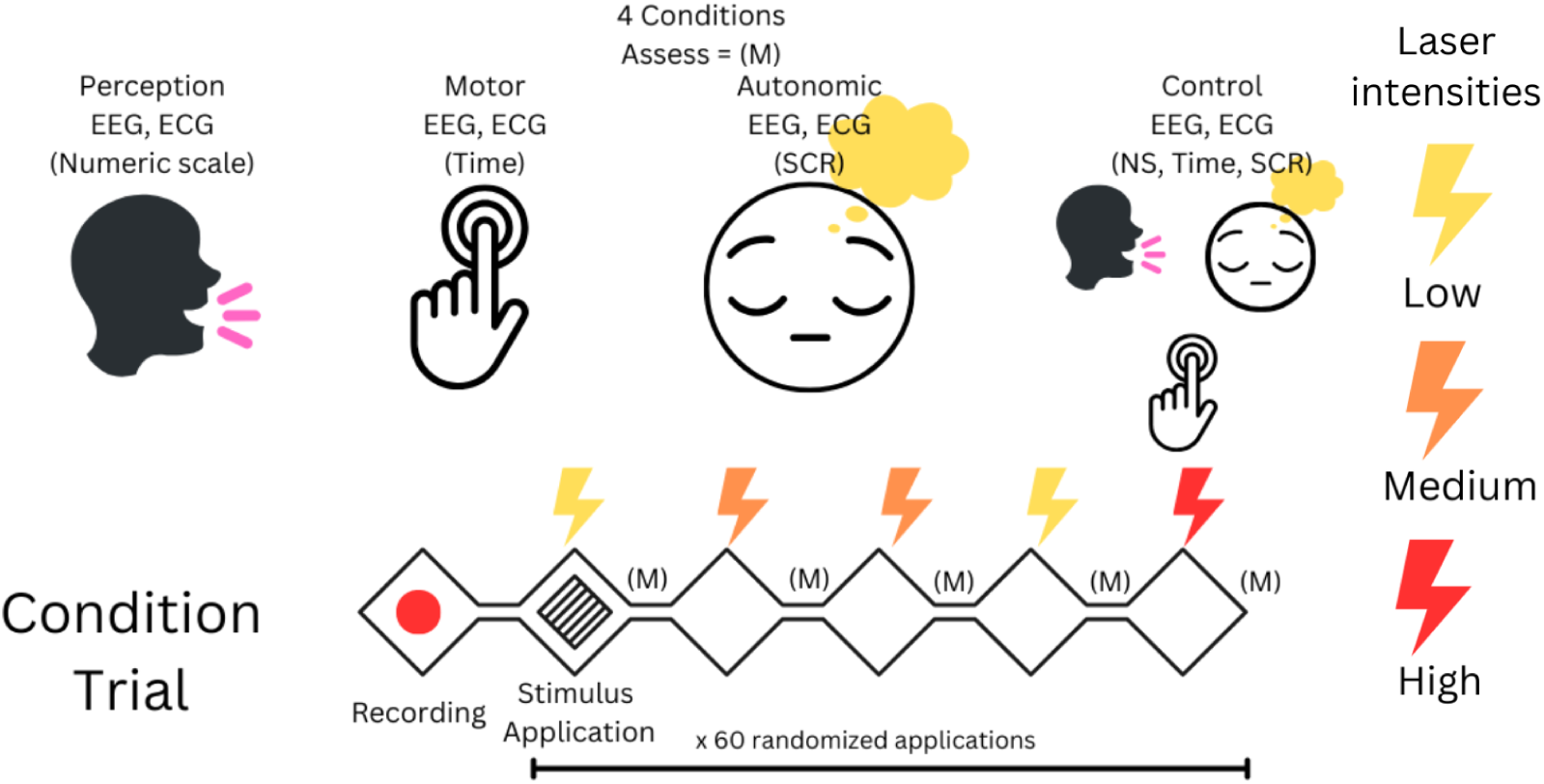
Experimental protocol. Four experiments were applied to each subject; Each experiment consisted of 60 trials (laser applications).

The perception condition consisted of the participant having to verbally evaluate the perception of pain on a scale from 0 to 100 after each laser application. Pain ratings serve as a measure of the perceptual dimension of pain. In the case of the motor condition, participants had to release a button that they were pressing as quickly as possible when the pain appeared; here, reaction times serve as a metric to measure the motor component of pain. In the autonomic condition, participants focused on the pain sensation without any other task, while the skin conductance response was recorded (SCR). This serves to measure the autonomic response of the body to pain. In the combined condition, participants first had to release the button as quickly as possible and then evaluate the pain from 1 to 100 on the pain rating scale, all while the EEG, ECG, and SCR were recorded. Subjects were instructed to keep their eyes closed during the experiments. For this research, only information on EEG signals is used.

### Data Acquisition

EEG data were recorded with an electrode cap (EasyCap, Herrsching) and BrainAmp MR plus amplifiers (Brain Products, Munich, Germany) using the BrainVision Recorder software (Brain Products, Munich, Germany). The electrode montage included 65 scalp electrodes consisting of all electrodes of the International 10–20 system as well as the additional electrodes FPz, AFz, FCz, CPz, POz, Oz, Iz, AF3/4, F5/6, FC1/2/3/4/5/6, FT7/8/9/10, C1/2/5/6, CP1/2/3/4/5/6, P1/2/5/6, TP7/8/9/10, and PO3/4/7/8/9/10.

Two additional electrodes were fixed below the outer canthus of each eye. This locations can be seen in the Fig. 3 During the recording, the EEG was referenced to the FCz electrode, grounded at AFz, sampled at 1000 Hz, highpass filtered at 0.015 Hz, and low-pass filtered at 250 Hz. The database also has a channel that contains information obtained from an electrocardiogram (ECG), which gives data on individuals’ heart rhythms. Finally, there is a channel for the sensor that measures the skin conductance response (SCR). It is worth mentioning that of the 65 EEG channels recorded by Tiemann et al.; in this research, only 62 were used. From the 62 channels used, 20 corresponded to the somatosensory cortex, since this area is linked to sensory and pain processes.

**Fig 3.**
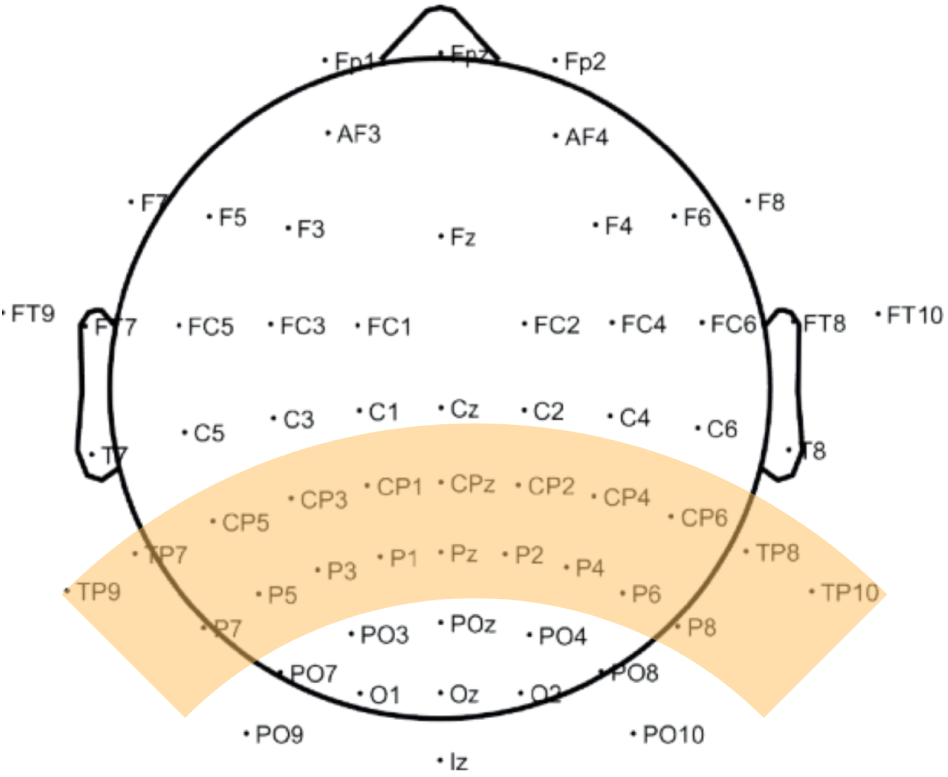
Location of the 62 electrodes used to record the dataset of nociceptive pain. In orange are placed the 20 channels of the somatosensory cortex.

## Methodology

This research focuses on detecting the change between the electrical signal at the moment of pain and the pre-stimulus, to know if there are different patterns between imagining that a noxious stimulus is coming and when it is already applied and finding a difference between different levels of pain that we assume exist when three stimuli are applied. This will be focused on certain characteristics in the frequency domain such as the power of specific frequency bands. The process for obtaining this data will be explained below.

### Pre-processing

MATLAB R2023a and EEGLAB [13] were used to pre-process the EEG signal and extract the features. As a first step, the data was down-sampled from 1000 Hz to 500 Hz; in this range, all oscillations of interest are possibly detectable in compliance with the laws of Nyquist, Theta (4.1-8 Hz), Alpha (8.1-12 Hz), Beta Low (12.5 - 20 Hz), Beta High (20.1 - 31 Hz), and Gamma (31.1 - 60 Hz). This is done to have less data and save time while it is processed. Then, a Notch filter of 50 Hz was used to eliminate electric line noise.

The EEG database is processed in this research with other biosignals (ECG) to apply independent component analysis (ICA) and more easily remove heart rate-related noise. In addition, electrodes placed under the eyelid will eliminate flicker-related noise. After this, the power of each band is obtained from the frequencies mentioned before.

**Fig 4.**
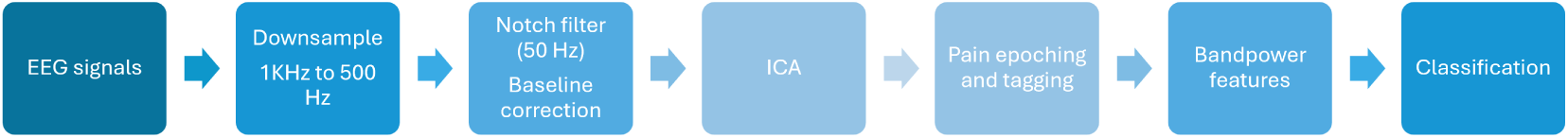
Timeline of the methodology used to preprocess the data.

### Feature Extraction and Classification

#### Bandpower

The features of interest in this study are the powers of the frequency bands because these are values that can show how much activity exists in a range of frequencies during a specific task. In this case, during the pain window.

The power of a frequency band in EEG is calculated by transforming the signal from the time domain to the frequency domain using the Fourier Transform. The power spectral density (PSD) was then obtained by squaring the magnitude of the frequency-domain signal. To isolate a specific band (e.g., alpha, beta), a band-pass filter is applied, and the PSD is integrated over the desired frequency range.

Fast Fourier transform (FFT):

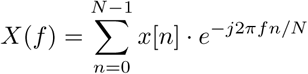

Power Spectral Density (PSD):

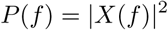

Band Power Integration:

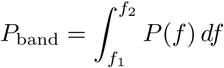

Where:

- *x*[*n*]: The EEG signal in the Time-domain.
- *X*(*f*): Frequency-domain representation of the signal.
- *P* (*f*): Power spectral density.
- *f*_1_ and *f*_2_: Lower and upper limits of the frequency band.

These windows are 1 second in length. In this case, 6 powers are extracted for each 1-second window. This is for each channel of the 62 used. So, for each of the 60 total, there will be 372 values related to the powers of the frequency bands (62 channels by 6 bands of frequency). In some cases, some laser applications (trials), have marking errors (such as when they missed the moment when the recording ended or when it started) or have recording errors so they have to be discarded.

#### Laser intensity labeling

As mentioned above, the way to generate pain was by applying a laser stimulus of 3 different intensities, High, Medium, and Low. Taking this into account, at the moment of calculating the powers of the frequency bands for each 1-second stimulus window, these were labeled depending on the stimulus applied. Thus, the labeling is dependent on the laser intensity. The stimulus would always be expected to cause a response of similar magnitude, and the subject’s perception is completely disregarded.

#### Time reaction labeling

For the time labeling, the subject’s perception is important. What is proposed here is that depending on the time of exposure to the stimulus, the level of pain for the labeling of the windows will be stimulated. The longer the reaction time, the pain is considered as low. If the reaction time is minimal, the pain is labeled as high. The maximum and minimum values function as a range, and the others are accommodated from highest to lowest between these. All are divided into 3 groups, having 20 values labeled as low pain, 20 as medium pain, and 20 as high pain.

**Fig 5.**
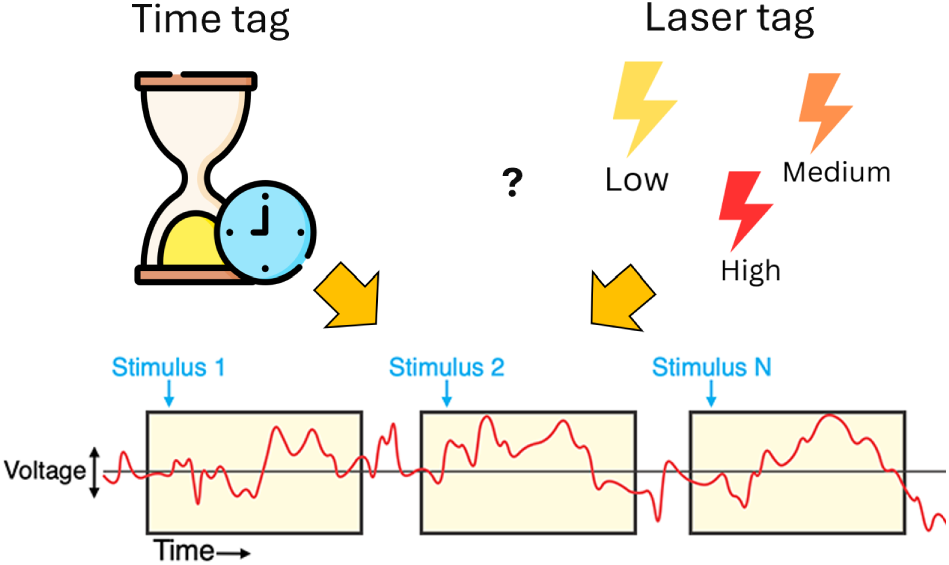
Time reaction tag and laser intensity tag feature labeling to each trial.

#### Classification

For classification, MATLAB’s Classification Learner was used, which has a series of algorithms that can be trained with data matrices. For this work the models were trained and tested per subject, using a 5-fold cross-validation. Therefore, 51 matrices with labels related to laser intensity and 51 related to the exposure time were used to obtain the results. The models used for this work were SVM (linear kernel), KNN (Euclidean distance to 1 neighbor), Neural Network (1 fully connected layer), and random forest. The results in this work are focused on the obtained using SVM, which has the best performance with this kind of data.

## Results

### Pain and Pre-stimulus

In this first analysis, the discrimination between the 1-second window of pain, and a 1-second window of the pre-stimulus (before the laser was applied) is of interest. This is to know if there is any change between the brain signal in its basal state and the brain signal at the moment of ”pain”. On the other hand, some research suggests that the expectation of a noxious stimulus generates a painful sensation similar to when this stimulus is already in contact with the body. The simple fact that we could face danger can generate an almost real imaginary pain [14–16].

The following results show a difference between pain and pre-pain (pre-stimulus), which is considered the moment when imaginary pain is present in the recorded participants.

The results showed were obtained using a linear Support Vector Machine (SVM) algorithm. This was the one that performed best when training and testing the data on a subject-by-subject basis. Previous work demonstrates the difference in algorithms between certain subjects in this database [17]. Figure 6 shows that the discrimination between pain and pre-stimulus states averages 85.72% using band power information from all channels (blue), and 81.94% using band power from the 20 channels located in the somatosensory cortex (orange). Using a greater number of channels compared to using only the 20 related to the somatosensory area seems to be better in most cases. When comparing the accuracies of two binary classification methods, a significant difference was found (p-value < 0.0000002) using a Student’s T-test, indicating that using more channels is statistically superior to the other in accuracy.

**Fig 6.**
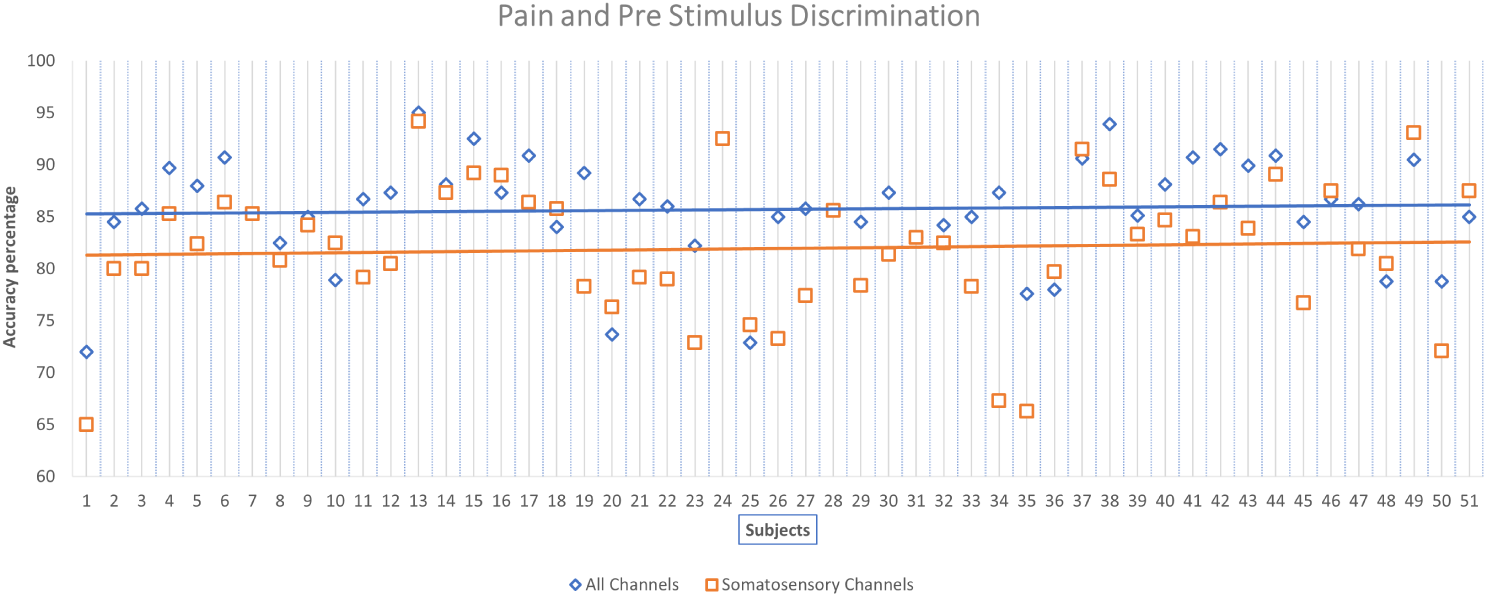
Accuracies from the classification using Linear SVM algorithm, blue is the accuracy of the data extracted from all the channels (62) and orange is the accuracy of the data extracted from the somatosensory channels (20).

### Low pain and High pain quantification

Given that there is activity in the pain window, different from the pre-stimulus, we were interested in exploring if the information in that window can be used to quantify it in levels. In this case, a binary discrimination was performed between the 1-second windows previously used but now labeled as Low Pain and High Pain windows.

The results obtained using the two labeling methods mentioned in the methodology will be shown below. The method called Laser Tag is completely related to the laser intensity that was applied to the subject and is taken directly from the markers used in the recording, which include this information at the beginning of each trial. Since the experiment consists of applying the laser 60 times −20 times for each intensity-only 20 of low pain and 20 of high pain were used.

In the case of Time Reaction Label, it is related to the time of exposure to the laser. It is taken from the moment when the laser was applied until the subject releases a button which indicates that they felt pain and the laser is deactivated. These times are ordered from highest to lowest and divided into 3 groups, trying to have 20 windows for each intensity. The first group, with higher values of time, is categorized as low pain since it means that the person “endured” the laser more depending on his perception of that stimulus.

The third group, with the shorter duration times, is labeled as High Pain, since we assume that the perception of those stimuli was so great that the subject did not endure so much exposure to them. The second group is labeled as medium and the values are milliseconds away from the other 2 groups, so they remain in this “limbo” due to the way the groups were divided. This will be discussed later. However, it is highly related to the way of quantifying between the three levels of pain.

Figure 7 shows the box plots of 4 cases, data labeled High and Low Laser Intensity Tags, Somatosensory Zone Laser Intensity Tags, Time Reaction Tags, and Somatosensory Zone Time Reaction Tags. As can be seen, the box with the best positioning is that of the classification model using the information of all channels with the labels related to the exposure time. This way of labeling achieved to have more subjects with a discrimination of two pain levels higher than 70 (30%). When compared with the box plot of the labeling concerning the laser, there is a 10% higher median and concentration of data when using labels related to exposure time and it is known that there are significant differences between both labels (p < 0.000017). In the case of using only the somatosensory zone channels in laser intensity labeling, the behavior appears to be similar to that of using all channels and there are no significant differences in the use of more or less data. The discrimination of the labeled data concerning time and using only 20 channels shows the highest dispersion of the 4 cases.

**Fig 7.**
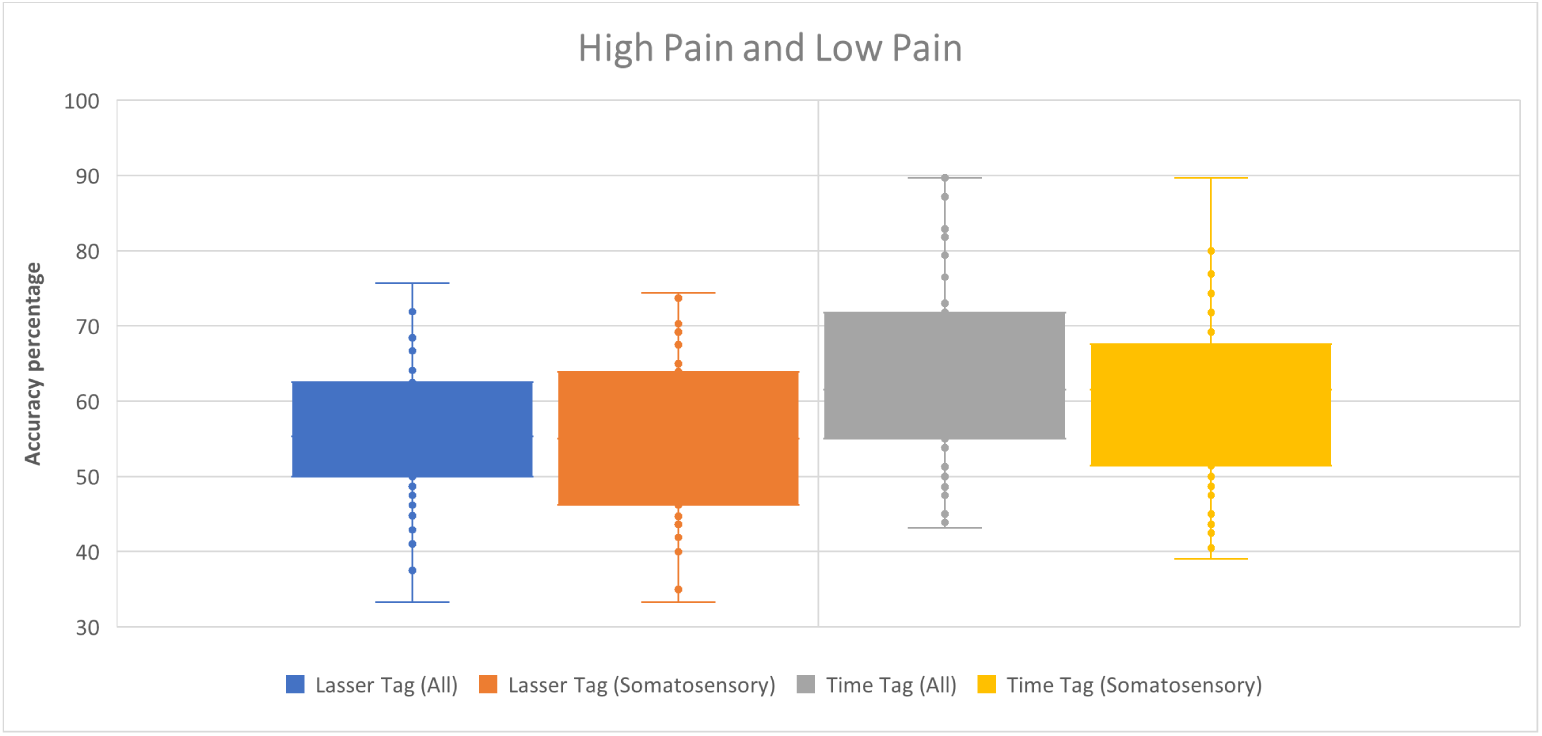
Boxplots from the accuracies of the binary discrimination between Low and High Pain using Linear SVM algorithm.

### High Pain, Medium Pain and Low Pain quantification

For the latter experiment, discrimination has been made between 3 levels of pain. The data labeled Medium Pain have been added. As mentioned in the previous section, in the case of laser-labeled data, these 20 new labels are taken by the intensity of the stimulus used. In the case of the time-related labels, the label is taken by the median of the group, the remaining 20 windows between the data labeled Low Pain and High Pain.

In the previous section, it is mentioned that this median pain, when labeled concerning time, is like the quartile left in “limbo” to say it colloquially. By this, we mean that many of the windows there could have belonged to high pain or low pain for milliseconds. And the average time that a subject lasts with exposure to pain is 0.3 seconds. This makes it more challenging to label what type of pain it was, but since they are quartiles, they have to stay in between, but the signals can have very similar patterns with the other 2 labeled.

In this last experiment, where the aim is to classify 3 levels of pain, the accuracies go down and reach between 35% and 40% on average when using laser intensity and time reaction labels, respectively as shown in Figure 8. It seems that the time reaction labels are better in general, compared to the laser intensity labels, but still have values that are negligible for tests where the aim is to quantify pain. The fact that using fewer channels shows a behavior similar to the previous cases, decreasing accuracies in most subjects. On the other hand, there are few and rare cases in which some of the subjects exceed 50% accuracy.

**Fig 8.**
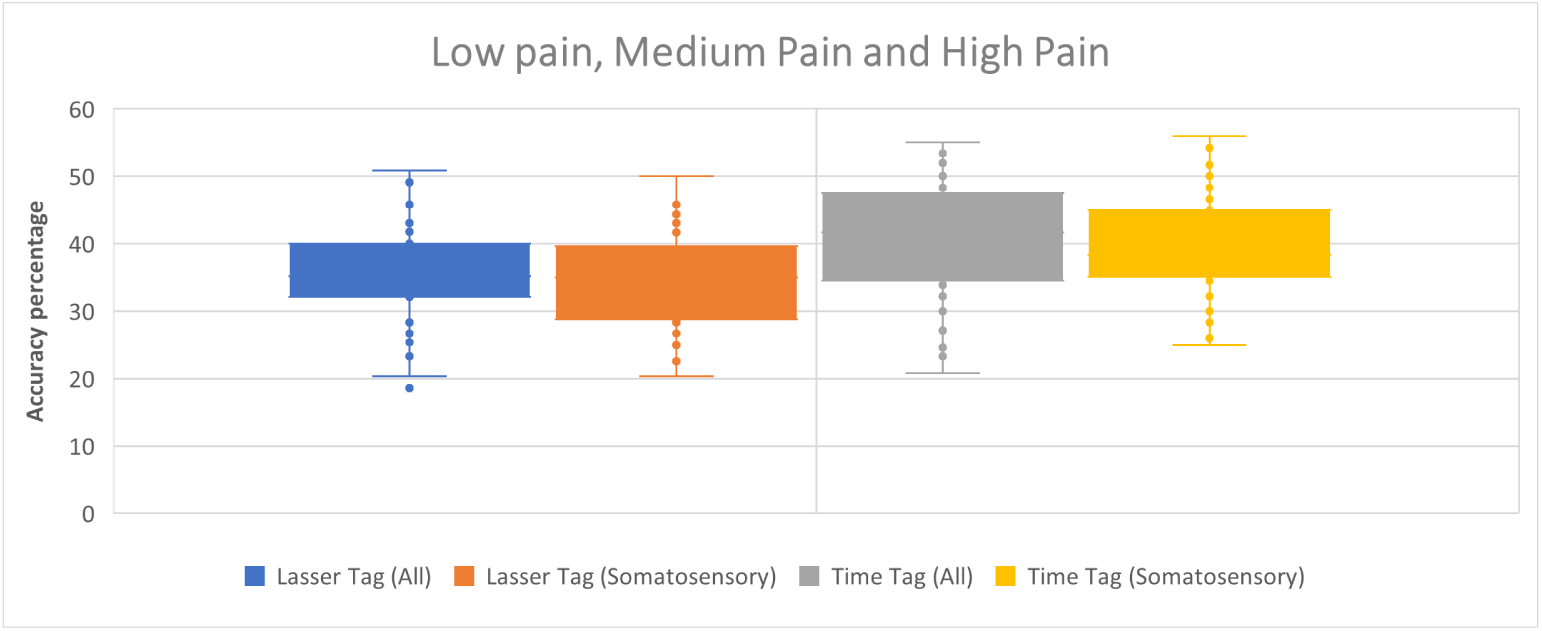
Boxplots from the accuracies of the discrimination between Low, Medium and High Pain using Linear SVM algorithm.

## Discussion

In this work has been demonstrated that it is possible to distinguish between states of pain (general) and pre-stimulus (pre-pain) with 90% accuracy in most subjects. This may be because an ON/OFF type phenomenon is generated in the brain signal when there is a stimulus and it is possible to know that it is different from the basal state of the subject, but the important thing is that it can be corroborated that the imaginary pain that is usually mentioned in certain articles about the fact of knowing that something will “hurt” us is very different from the real harmful stimulus already in contact. It also means that in the 1-second windows taken to extract features, there is activity that may be useful for the next two experiments.

In images 9 and 10 it can be observed how the labeling of the signals influences an evident change of the pain window and the pre-stimulus window, supporting the previous results. In the case of labeling concerning stimulus intensity, the peaks at the 100 positive milliseconds (P100) are more evident compared to labeling concerning reaction time. But, in the case of labeling for reaction time, a greater difference can be observed in the power of the high signal milliseconds after 100 milliseconds. On the other hand, in the labeling related to laser intensity, there seems to be a similarity between low and medium intensity. While in the reaction time labeling, all the intensities coincide in power in the first positive peak, but during the pain window, they are quite different from each other.

In the case of low pain and high pain, it is observed that approximately 30% of the subjects manage to obtain more than 70% accuracy when discriminating with labels related to the exposure time. This was the method that obtained more interest since it shows indications that there are differences in the powers of the frequency bands when the pain stimulus changes in its intensity. For the third experiment, we sought to add one more level, the medium level. But here the learning models simply worked randomly in almost all cases, performing very poorly in discriminating between 3 levels of pain.

**Fig 9.**
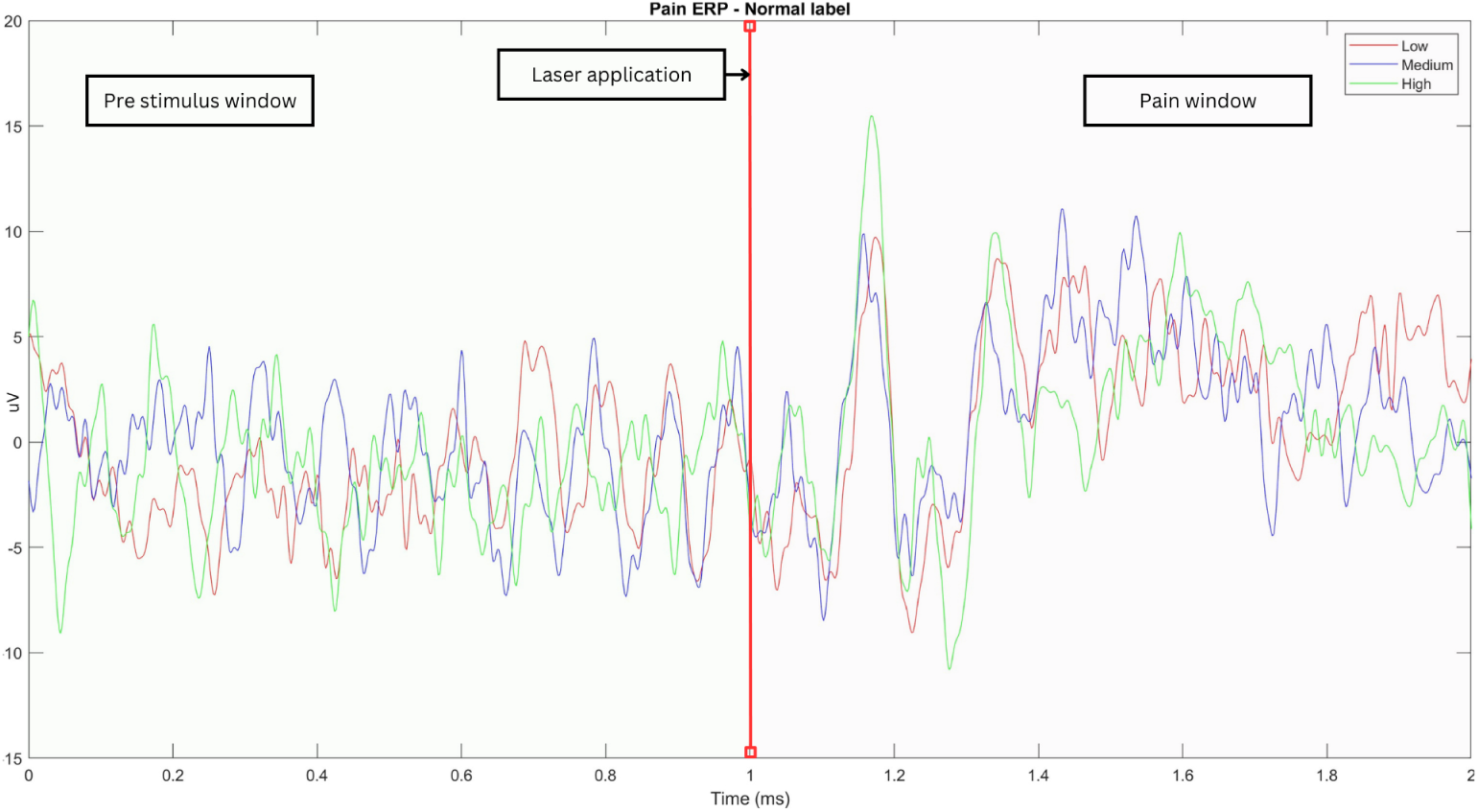
Window of pre-stimulus and pain from channel CPz Event-Related Potential (ERP) of subject 13 when the data is labeled as Laser intensity tagging.

**Fig 10.**
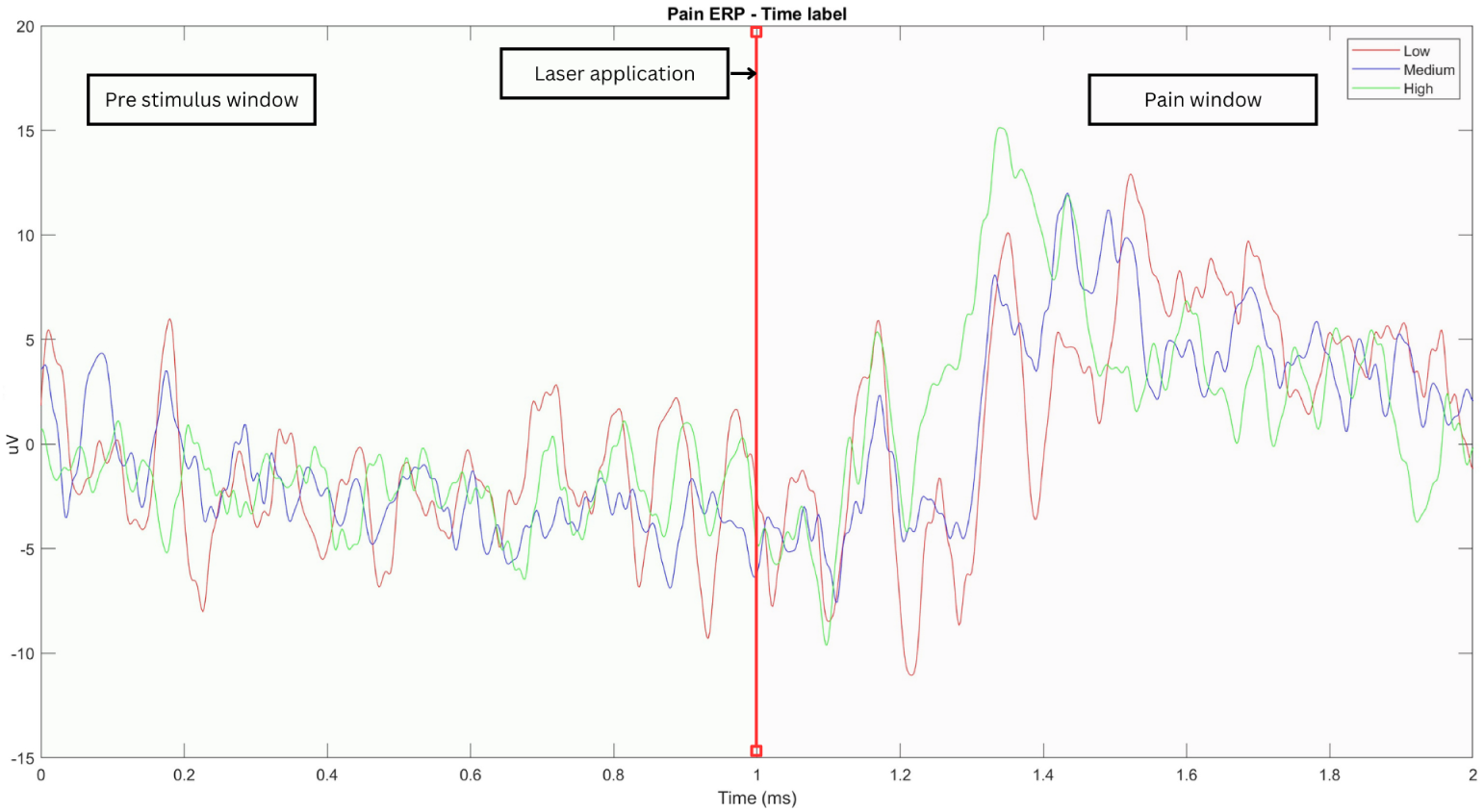
Window of pre-stimulus and pain from channel CPz Event-Related Potential (ERP) of subject 13 when the data is labeled as Time reaction tagging.

Even using the time reaction labeling method, it is usually very poor performance, as shown in Figure 11. But here arises one of the most important questions in this research.

**Fig 11.**
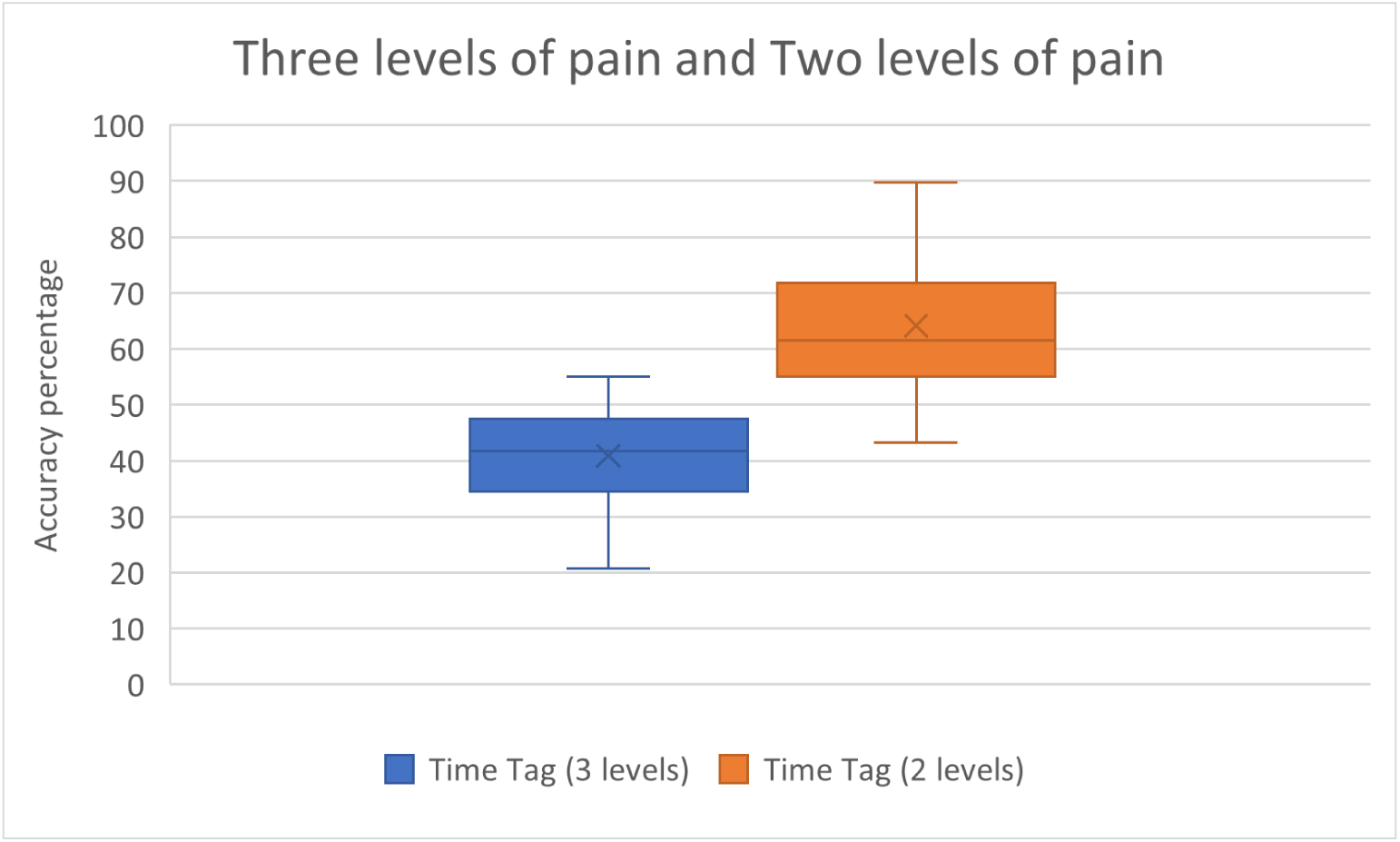
Boxplots from the accuracies of the discrimination between the quantification of 3 levels of pain vs de 2 levels of pain tagged with time information.

What would be the best way to determine whether a pain is high, medium or low?

It seems that taking into account the temporal information of the reaction is what worked best in this research. But by creating groups, which are arbitrarily generated by dividing a list of values ordered in descending order into equal parts, values very similar to those of the groups at the extremes (low and high) and which were left in the “medium” group, create confusion in the classification models. When perhaps in the end it is very similar in power and time to some data of the other two groups. This gives rise to plan and think the best way to label something that is completely subjective, and perhaps to implement some other model of labeling in conjunction with the time, to define these levels.

On the other hand, the scales used in data collection are scales that go from 0 to 100, and we are trying to create 3 groups. So, what if we had to come up with a quantification model where the final label is not high, medium and low, but a numerical scale of 101 digits? This would seem to increase the difficulty, but it may have to do with the perception and judgment that the person will have at the time of giving their rating to the pain experienced.

That said, the results shown here are evidence that there are possibilities of detecting pain and quantifying it in 2 levels for the moment.

## Conclusion

The field of pain quantification with bio-signals continues to grow, and what is documented here are steps toward a better understanding of this type of subjective experience. A better method of labeling the signals needs to be evaluated and designed so that typical learning models can discriminate between pain levels and perhaps even give a number on a scale of certain values.

For the time being, we will continue to work with these data and we have also started to work on classifying the time-frequency images of these pain windows. To know if using image classification methods it is possible to differentiate between images and quantify pain levels.

It would also be interesting to know if there is an index or a value in the frequency bands or the information of the electrical signal with which labeling can be given, so the analysis of EEG related to pain is also in work.

The authors of this corpus, mentions certain patterns that can be useful to understand the signal generated by pain, perhaps the amplitudes, desynchronizations or synchronizations and some more data related to the nature of the brain signal give clues to achieve a more successful quantification of nociceptive pain.

For now, this work shows that the use of frequency characteristics such as band power is useful for classifying pain intensities. The type of labeling plays an important role in generating the data and by taking into account the subjects’ perception (time of reaction), it seems to be an effective way to solve this problem.

## Supporting information

Link for the tables of accuracies of all the subjects ordered into types of labeling and with information on the different classification models used.

## Acknowledgments

We thank INAOE for the academic and financial support provided throughout this research. To the University of Winnipeg for the collaboration and funding of this article. And to the SECIHTI for the postgraduate grant (CVU 908456). Also, to Tiemann et al. for the useful dataset collected.

